## Supplemental File 3 for "Quantitative proteomics reveals key roles for post-transcriptional gene regulation in the molecular pathology of FSHD": Supp-file-3_proteomics-analysis.Rmd

````
---
title: "DUX4_proteomics"
author: "Sujatha Jagannathan"
date: "June 7, 2018"
output: 
  html_document: 
    toc: true
    toc_float: true
    toc_collapsed: false
    fig_caption: yes
    fig_retina: 1 
    df_print: paged
---
###Instructions to use this RMarkdown document
1. Place the .Rmd document in any folder and set that folder as the working directory
2. Create folders as follows
    <current-dir> > data > mass_spec
    <current-dir> > data > rdata
3. The files needed to knit this document are:
    Four psm files placed within data/mass_spec
    Uniprot sequences placed within data/rdata
    RNAseq data placed within data/rdata
4. With all of the above in place, the document should knit and output a html files with several figure panels as well as create many files containing intermediate data and the supplementary files 1 and 2. 

###Set up
Required packages: dplyr, tidyr, stringr, ggplot2
```{r}
library(shiny)
library(tidyverse)
library(ggplot2)
library(plotly)
library(gridExtra)
set.seed(66958473)

setwd("/Users/sjaganna/Dropbox/Manuscripts/2018_Jagannathan_etal_DUX4_proteomics")
```

###Enter data: 
Use tab-delimited PSM files from Mass spec core for each of the replicates
```{r, psmdata_input}
library(readr)
psmdata_vDUX4_24h_rep0 <- read_delim("data/mass_spec/SJ102314_102914_05_16_TargetPeptideSpectrumMatch.txt", "\t", escape_double = FALSE, trim_ws = TRUE)

psmdata_vDUX4_36h_rep0 <- read_delim("data/mass_spec/SJ102314_102914_17_28_TargetPeptideSpectrumMatch.txt", "\t", escape_double = FALSE, trim_ws = TRUE)

psmdata_iDUX4_14h_rep1 <- read_delim("data/mass_spec/SJ042415_051615_31_46_TargetPeptideSpectrumMatch.txt", "\t", escape_double = FALSE, trim_ws = TRUE)

psmdata_iDUX4_14h_rep2 <- read_delim("data/mass_spec/SJ090815_091515_47_62_TargetPeptideSpectrumMatch.txt", "\t", escape_double = FALSE, trim_ws = TRUE)

psmdata_iDUX4_14h_rep1 <- dplyr::tbl_df(psmdata_iDUX4_14h_rep1)
psmdata_iDUX4_14h_rep2 <- dplyr::tbl_df(psmdata_iDUX4_14h_rep2)
psmdata_vDUX4_24h_rep0 <- dplyr::tbl_df(psmdata_vDUX4_24h_rep0)
psmdata_vDUX4_36h_rep0 <- dplyr::tbl_df(psmdata_vDUX4_36h_rep0)
```


1.Giving meaningful names to the columns to enable subsetting as follows: 
```{r}
colnames.vDUX4 = c("Annotated Sequence"="peptide.sequence", "Modifications"="modifications", "Master Protein Accessions"="masterids", "Protein Accessions"="ids", "# Protein Groups"="num.masterids", "# Proteins"="num.ids", "# Missing Channels"="missing.channel", "# Single-Peak Channels"="single.peak", "Percolator q-Value"="q.value", "Percolator PEP"="pep.value", "Quan Info"="quan.info", "Peptide Quan Usage"="quan.usage", "Precursor Quan Result ID"="quan.id", "Spectrum File"="band.id", "Area"="area", "Light"="ctrl", "Heavy"="dux4")

colnames.iDUX4.rep1 = c("Annotated Sequence"="peptide.sequence", "Modifications"="modifications", "Master Protein Accessions"="masterids", "Protein Accessions"="ids", "# Protein Groups"="num.masterids", "# Proteins"="num.ids", "# Missing Channels"="missing.channel", "# Single-Peak Channels"="single.peak", "Percolator q-Value"="q.value", "Percolator PEP"="pep.value", "Quan Info"="quan.info", "Peptide Quan Usage"="quan.usage", "Precursor Quan Result ID"="quan.id", "Spectrum File"="band.id", "Area"="area", "Light"="ctrl", "Heavy"="dux4")

#label swap replicate
colnames.iDUX4.rep2 = c("Annotated Sequence"="peptide.sequence", "Modifications"="modifications", "Master Protein Accessions"="masterids", "Protein Accessions"="ids", "# Protein Groups"="num.masterids", "# Proteins"="num.ids", "# Missing Channels"="missing.channel", "# Single-Peak Channels"="single.peak", "Percolator q-Value"="q.value", "Percolator PEP"="pep.value", "Quan Info"="quan.info", "Peptide Quan Usage"="quan.usage", "Precursor Quan Result ID"="quan.id", "Spectrum File"="band.id", "Area"="area", "Light"="dux4", "Heavy"="ctrl")

psmdata_vDUX4_24h_rep0 = psmdata_vDUX4_24h_rep0[, names(colnames.vDUX4)]
colnames(psmdata_vDUX4_24h_rep0)=colnames.vDUX4

psmdata_vDUX4_36h_rep0 = psmdata_vDUX4_36h_rep0[, names(colnames.vDUX4)]
colnames(psmdata_vDUX4_36h_rep0)=colnames.vDUX4

psmdata_iDUX4_14h_rep1 = psmdata_iDUX4_14h_rep1[, names(colnames.iDUX4.rep1)]
colnames(psmdata_iDUX4_14h_rep1)=colnames.iDUX4.rep1

psmdata_iDUX4_14h_rep2 = psmdata_iDUX4_14h_rep2[, names(colnames.iDUX4.rep2)]
colnames(psmdata_iDUX4_14h_rep2)=colnames.iDUX4.rep2

library(knitr)
kable(cbind(psms_iDUX4_rep1 = nrow(psmdata_iDUX4_14h_rep1), 
            psms_iDUX4_rep2 = nrow(psmdata_iDUX4_14h_rep2), 
            psms_vDUX4_24h = nrow(psmdata_vDUX4_24h_rep0), 
            psms_vDUX4_36h = nrow(psmdata_vDUX4_36h_rep0)))

```  


2. Merge the two reps after assigning "rep" id
```{r}
psmdata_vDUX4_24h_rep0 <- psmdata_vDUX4_24h_rep0 %>% mutate(rep = "1")
psmdata_vDUX4_36h_rep0 <- psmdata_vDUX4_36h_rep0 %>% mutate(rep = "2")
psmdata_iDUX4_14h_rep1 <- psmdata_iDUX4_14h_rep1 %>% mutate(rep = "3")
psmdata_iDUX4_14h_rep2 <- psmdata_iDUX4_14h_rep2 %>% mutate(rep = "4")
psmdata.static <- rbind(psmdata_vDUX4_24h_rep0, psmdata_vDUX4_36h_rep0, psmdata_iDUX4_14h_rep1, psmdata_iDUX4_14h_rep2)
```  


3. assigning peptide ids as row names for each of the `r nrow(psmdata.static)` rows  
```{r}
row.names = paste0("peptide", c(1:nrow(psmdata.static)))
rownames(psmdata.static)=row.names
```  


4. filter psmdata to remove all but unique peptides and exlcudedbymethod peptides (which includes missing.channel psms in the case of replicate 1):  
```{r}
library(knitr)
kable(cbind(psms = table(psmdata.static$quan.info))) 
psmdata.static.filtered <- psmdata.static[which(psmdata.static$quan.info %in% c("Unique", "ExcludedByMethod")), ]
```
Before filtering: `r kable(cbind(psms = table(psmdata.static$quan.info)))`  
After filtering: `r kable(cbind(psms = table(psmdata.static.filtered$quan.info)))`  
Of the filtered psms, there are `r table(psmdata.static.filtered$single.peak)` psms with single peaks and `r table(psmdata.static.filtered$missing.channel)` psms with missing channels.   

5. Clean up data
  a) assign single.peak and missing.channel
  b) convert peptide sequence to upper case for string matching
  c) recalculate hl.ratio to account for iDUX4_rep2 having labels swapped 
  d) filter our rows with no masterid assigned
```{r}
library("stringr")
psmdata.static.filtered <- psmdata.static.filtered %>%
  mutate(quan.info=ifelse(is.na(missing.channel), quan.info, "missing.channel")) %>%
  mutate(quan.info=ifelse(is.na(single.peak), quan.info, "single.peak")) %>% 
  mutate(peptide.sequence=str_to_upper(peptide.sequence)) %>%
  mutate(hl.ratio = dux4/ctrl) %>%
  filter(masterids != "")
```  
The resulting psmdata file has the following psms: `r kable(cbind(psms = table(psmdata.static.filtered$quan.info)))`

6. Drop the missing channel and single peak columns as that information is now in the quan.info column  
```{r}
psmdata.static.filtered <- psmdata.static.filtered[, c(1:6, 9:19)]
```  

7. Filter out single.peak and ExcludedByMethod psms  
```{r}
psmdata.static.filtered <- psmdata.static.filtered %>%
                      filter(quan.info != "single.peak") %>% 
                        filter(quan.info != "ExcludedByMethod")
```  

8. (Optional) Export 'psmdata' with `r nrow(psmdata.static.filtered)` psms as an R object  
```{r}
save(psmdata.static.filtered, file="data/rdata/psmdata.06072018.Rdata")  
```  

###Data transformation  
1. Remove psms that map to contaminants from cRAP database (all of them have the id format of 'sp')
```{r}
psmdata.static.filtered <- psmdata.static.filtered %>% filter(!grepl("; sp|^sp", ids))
```  
This step leaves us with `r nrow(psmdata.static.filtered)` psms.  

2. Replace the low missing value with 10000 to make it less fluctuating (and to not have 2 separate groups of points near the edges of the rep1 vs rep2 scatter plot) and recalculate hl.ratio and log.hl.ratio  
```{r}
psmdata.static.filtered <- psmdata.static.filtered %>% mutate(ctrl = ifelse(ctrl < 10000, 10000, ctrl)) %>% 
                      mutate(dux4 = ifelse(dux4 < 10000, 10000, dux4)) %>% 
                          mutate(hl.ratio = dux4/ctrl, log.hl.ratio = log2(dux4/ctrl))
```   

3. Normalize using means for rep1 and rep2 separately  
```{r}
tmp <- psmdata.static.filtered[-which(psmdata.static.filtered$quan.info %in% c("missing.channel")), ]
rep1 <- tmp[which(tmp$rep=="1"), ]
rep2 <- tmp[which(tmp$rep=="2"), ]
rep3 <- tmp[which(tmp$rep=="3"), ]
rep4 <- tmp[which(tmp$rep=="4"), ]

rep1.mean <- mean(rep1$log.hl.ratio)
rep2.mean <- mean(rep2$log.hl.ratio)
rep3.mean <- mean(rep3$log.hl.ratio)
rep4.mean <- mean(rep4$log.hl.ratio)
rep1.sd <- sd(rep1$log.hl.ratio)
rep2.sd <- sd(rep2$log.hl.ratio)
rep3.sd <- sd(rep3$log.hl.ratio)
rep4.sd <- sd(rep4$log.hl.ratio)

psmdata.static.norm <- psmdata.static.filtered %>% 
                  mutate(norm.hl.ratio = ifelse(rep=="1", (log.hl.ratio-rep1.mean)/rep1.sd, 
                                                ifelse(rep=="2", (log.hl.ratio-rep2.mean)/rep2.sd, 
                                                       ifelse(rep=="3", (log.hl.ratio-rep3.mean)/rep3.sd, 
                                                              (log.hl.ratio-rep4.mean)/rep4.sd))))

psmdata.static.norm <- psmdata.static.norm %>% mutate(peptide.sequence = str_replace_all(psmdata.static.norm$peptide.sequence, "[[:punct:]]", ""))
```  
`r kable(cbind(Rep1_mean = rep1.mean, Rep2_mean = rep2.mean, Rep1_sd = rep1.sd, Rep2_sd = rep2.sd))`

###Data summarization  
1. Creating a new datastructure that maps geneid -> peptide: In this step, the data structure shifts from one-peptide/many-ids to several one-peptide/one-id rows. I can then summarize these data after grouping by geneid (i. e. one-id/many-peptides). Since I am limiting myself to only unique peptides, most of my psms are one-peptide/one-id anyway. So this step is not as important as it used to be when I was not filtering out the NotUnique peptides, but I do it anyway (Note: some psms are assigned to multiple masterids, but are not listed as NotUnique and were hence not filtered out. How to handle this? GCC2 is one example)
```{r}
library("stringr")

psmdata.static.masterid <- psmdata.static.norm %>%
  # Figure out how many geneids the peptide mapped to
  mutate(nmasterids=str_count(psmdata.static.norm$masterids, pattern = ";") + 1, 
         N = row_number()) %>% 
  # Create a data frame with one geneid per row
  do(data.frame(.[rep(.$N, .$nmasterids), ], 
                masterid=unlist(strsplit(.$masterids, "; ", fixed=TRUE))))
```

2. Mapping uniprot ids to geneids and filtering out uncharacterized genes
```{r}
##Importing uniprot id -> geneid map
load("data/rdata/uniprot.ids.sequences.Rdata")

##Joining the geneid information to psmdata dataframes and filtering uncharacterized and duplicate entries

psmdata.geneid.static <- left_join(psmdata.static.masterid, uniprot.ids.sequences, by = "masterid")
psmdata.geneid.static <- psmdata.geneid.static[, c(22:24, 19, 1:18, 25)]
psmdata.geneid.static <- filter(psmdata.geneid.static, geneid!="Uncharacterized")

```


3. printing the position of peptide in the protein (to allow peptide position mapping later)
```{r}
library("stringr")
match_position <- as.data.frame(str_locate(as.character(psmdata.geneid.static$sequences), psmdata.geneid.static$peptide.sequence))
protein.length <- str_length(as.character(psmdata.geneid.static$sequences))
psmdata.geneid.static <- cbind(psmdata.geneid.static, peptide.position = match_position$start, protein.length)
psmdata.geneid.static <- psmdata.geneid.static[, c(1:22, 24:25, 23)]
psmdata.geneid.static <- psmdata.geneid.static %>% 
                      mutate (peptide.position.index = peptide.position/protein.length)
psmdata.geneid.static <- psmdata.geneid.static[, c(1:24, 26, 25)]
```  

4. (optional) export the data
```{r}
save(psmdata.geneid.static, file="data/rdata/psmdata.geneid.static.06072018.Rdata")
```

###Bootstrapping to get significance  
1. Generate a table containing peptide.sequence, masterids and norm.hl.ratio
2. Shuffle the norm.hl.ratios randomly using sample() and do this 1000 times (should I do more?)
3. Proceed to summarization of the peptide hl.ratios to get protein ratios (1000 values for each protein obtained by taking the median hl.ratio of all the peptides)- the bootstrap table will provide a distribution of protein hl.ratios derived from randomly assigned peptide ratios. Hence, the mean and standard deviation will depend on how many peptides were summarized into the protein hl.ratio. Fewer peptides would mean more noise. From this table, I can calculate the mean and sd of protein hl.ratios expected under the null hypothesis and use this to derive a p.value for the actual observed ratio. 
```{r}
psmdata.static.norm.bootstrap <- psmdata.static.norm %>% filter(rep != "1", rep!= "2")
bootstrap <- data.frame(cbind(psmdata.static.norm.bootstrap[, c(1, 3, 19)], replicate(1000, {sample(psmdata.static.norm.bootstrap$norm.hl.ratio)})))

#bootstrap data
bootstrap.masterid <- bootstrap %>% 
   # Figure out how many geneids the peptide mapped to
  mutate(nmasterids=str_count(bootstrap$masterids, pattern = ";") + 1, 
         N = row_number()) %>% 
  # Create a data frame with one geneid per row
  do(data.frame(.[rep(.$N, .$nmasterids), ], 
                masterid=unlist(strsplit(.$masterids, "; ", fixed=TRUE))))

bootstrap.geneid <- left_join(bootstrap.masterid, uniprot.ids.sequences[, c(1:2)], by = "masterid")
bootstrap.geneid <- bootstrap.geneid[, c(1:3, 1006:1007, 4:1003)]
bootstrap.geneid <- filter(bootstrap.geneid, geneid!="Uncharacterized")
```

###Data Summarization
1. Calculating hl.ratio by median: for both both psmdata and bootstrap data. Other parameters calculated are: Inter Quartile Range (IQR), Quartile Coefficient of Deviation (QCoD) and peptide count. 
```{r}
psmdata.rep1stats <- psmdata.geneid.static %>%
  filter(rep=="1") %>%
  group_by(geneid) %>% 
  summarize (hl.ratio.rep1 = median(norm.hl.ratio), 
             area.control.rep1 = median(ctrl),
             area.dux4.rep1 = median(dux4),
             area.rep1 = median(area),
             IQR.rep1 = IQR(norm.hl.ratio, type=7), 
             QCoD.rep1 = IQR.rep1/hl.ratio.rep1, 
             count.rep1 = n())

psmdata.rep2stats <- psmdata.geneid.static %>% 
  filter(rep=="2") %>%
  group_by(geneid) %>% 
  summarize (hl.ratio.rep2 = median(norm.hl.ratio), 
             area.control.rep2 = median(ctrl),
             area.dux4.rep2 = median(dux4),
             area.rep2 = median(area),
             IQR.rep2 = IQR(norm.hl.ratio, type=7), 
             QCoD.rep2 = IQR.rep2/hl.ratio.rep2, 
             count.rep2 = n())

psmdata.rep3stats <- psmdata.geneid.static %>%
  filter(rep=="3") %>%
  group_by(geneid) %>% 
  summarize (hl.ratio.rep3 = median(norm.hl.ratio), 
             area.control.rep3 = median(ctrl),
             area.dux4.rep3 = median(dux4),
             area.rep3 = median(area),
             IQR.rep3 = IQR(norm.hl.ratio, type=7), 
             QCoD.rep3 = IQR.rep3/hl.ratio.rep3, 
             count.rep3 = n())

psmdata.rep4stats <- psmdata.geneid.static %>% 
  filter(rep=="4") %>%
  group_by(geneid) %>% 
  summarize (hl.ratio.rep4 = median(norm.hl.ratio), 
             area.control.rep4 = median(ctrl),
             area.dux4.rep4 = median(dux4),
             area.rep4 = median(area),
             IQR.rep4 = IQR(norm.hl.ratio, type=7), 
             QCoD.rep4 = IQR.rep4/hl.ratio.rep4, 
             count.rep4 = n())

psmdata.repstats <- psmdata.geneid.static %>% 
    filter(rep!= "1", rep!= "2") %>%
  group_by(geneid) %>% 
  summarize (hl.ratio=median(norm.hl.ratio), 
             area.control = median(ctrl),
             area.dux4 = median(dux4),
             area = median(area),
             protein.length = mean(protein.length), 
             rep=mean(as.numeric(rep)), 
             IQR = IQR(norm.hl.ratio, type=7), 
             QCoD = IQR/hl.ratio, 
             count = n())

psmdata.summary.static <- full_join(psmdata.rep1stats, psmdata.rep2stats, by="geneid")
psmdata.summary.static <- full_join(psmdata.summary.static, psmdata.rep3stats, by="geneid")
psmdata.summary.static <- full_join(psmdata.summary.static, psmdata.rep4stats, by="geneid")
psmdata.summary.static <- full_join(psmdata.summary.static, psmdata.repstats, by="geneid")

#bootstrap
bootstrap.summary <- bootstrap.geneid[, c(5:1005)] %>%
  group_by(geneid) %>% 
  summarise_all (funs(median))

bootstrap.summary.formeanandsd <- bootstrap.summary[, c(2:1001)]

bootstrap.mean.sd <- data.frame(cbind(mean = rowMeans(bootstrap.summary.formeanandsd), sd=apply(bootstrap.summary.formeanandsd,1, sd, na.rm = TRUE)))

bootstrap.mean.sd <- cbind(bootstrap.summary$geneid, bootstrap.mean.sd)

names(bootstrap.mean.sd) <- c("geneid", "mean", "sd")
```

2. Estimate p.value by using the bootstrap distribution of protein hl.ratios  
```{r}
psmdata.summary.pval.static <- full_join(psmdata.summary.static, bootstrap.mean.sd, by = "geneid")
psmdata.summary.pval.static <- psmdata.summary.pval.static %>% 
                          mutate(zscore = (hl.ratio-mean)/sd) 

pval <- 2*pnorm(-(abs(psmdata.summary.pval.static$zscore)))
psmdata.summary.pval.static <- cbind(psmdata.summary.pval.static, pval)

# psmdata.summary <- cbind(psmdata.summary, 
#                                p.value = 2*pnorm(-(abs(psmdata.summary$hl.ratio))))
# 
# nrow(psmdata.summary)
# #5240
# psmdata.summary <- cbind(psmdata.summary, 
#                                adjusted.p.value = p.adjust(
#                                  psmdata.summary$p.value, 
#                                  method="hochberg", 
#                                  n = 5245))
```  

3. Importing RNA-seq data from MB135 with vDUX4 and iDUX4 (already saved as an R object) - from the HMG paper analysis
```{r}
load("data/rdata/edgeR_foldchange_unfiltered.Rdata")
rnaseq.unfiltered <- foldchange.unfiltered[, c(2:7, 13:17)]
names(rnaseq.unfiltered) <- c("geneid", "iDUX4_logFC_UF", "iDUX4_logCPM_UF", "iDUX4_LR_UF", "iDUX4_pval_UF", "iDUX4_fdr_UF", "vDUX4_logFC_UF", "vDUX4_logCPM_UF", "vDUX4_LR_UF", "vDUX4_pval_UF", "vDUX4_fdr_UF")

library(readr)
rnaseq.filtered <- read_delim("data/rnaseq.filtered.txt", "\t", escape_double = FALSE, trim_ws = TRUE)
rnaseq.filtered <- rnaseq.filtered[, c(2:7, 13:17)]

names(rnaseq.filtered) <- c("geneid", "iDUX4_logFC", "iDUX4_logCPM", "iDUX4_LR", "iDUX4_pval", "iDUX4_fdr", "vDUX4_logFC", "vDUX4_logCPM", "vDUX4_LR", "vDUX4_pval", "vDUX4_fdr")

rnaseq <- full_join(rnaseq.filtered, rnaseq.unfiltered, by = "geneid")
``` 

4. Merging RNA-seq data with SILAC data using full_join
```{r}
data.rnaseq.silac.static <- full_join(rnaseq, psmdata.summary.pval.static, by = "geneid")
```
Final number of protein ids: `r nrow(data.rnaseq.silac.static)`

5. (Optional) Exporting data
```{r}
save(data.rnaseq.silac.static, file="data/rdata/data.rnaseq.silac.static_06072018.Rdata")
write.csv(data.rnaseq.silac.static, file="data/rdata/data.rnaseq.silac.static_06072018.csv")

```  

###Figures

0. Setting up plots
```{r}
library(tidyverse)
library(ggplot2)
library(plotly)
library(gridExtra)

mytheme <- theme_minimal(base_size = 15) + theme(aspect.ratio = 1, 
                                                 panel.grid.major = element_blank(),
                                                 panel.grid.minor = element_blank(),
                                                 panel.background = element_rect(colour = "black", size=1))
```

Figure 1 panels
```{r}
x <- psmdata.geneid.static$norm.hl.ratio
bins <- 40
       
# draw the histogram with the specified number of bins

##ZSCAN4
ggplot(psmdata.geneid.static %>% 
         filter(geneid == "ZSCAN4"), aes(norm.hl.ratio)) + 
         scale_x_continuous(limits=c(-20, 20)) +
         geom_histogram(bins = bins, fill = "#10101080") +
         mytheme +
         xlab("Normalized DUX4/Control ratio (log2)") +
         ylab("Peptide count") +
         facet_wrap(~rep, ncol =2, labeller = label_parsed) +
         geom_vline(xintercept = 0, linetype="dashed")

##RPL15
ggplot(psmdata.geneid.static %>% 
         filter(geneid == "RPL15"), aes(norm.hl.ratio)) + 
         scale_x_continuous(limits=c(-20, 20)) +
         geom_histogram(bins = bins, fill = "#10101080") +
         mytheme +
         xlab("Normalized DUX4/Control ratio (log2)") +
         ylab("Peptide count") +
         facet_wrap(~rep, ncol =2, labeller = label_parsed) +
         geom_vline(xintercept = 0, linetype="dashed")

# boxplots for yao biomarkers

# genes are from yao et al, table #3 - fshd biomarkers
yao.genes <- c("AC010606.1", "ALPPL2", "C1DP2", "CCNA1", "DUXA", "HNRNPCL1", "KDM4E", "KHDC1L", "KHDC1L", "KHDC1P1", "
KLF17", "LEUTX", "MBD3L2", "MBD3L3", "MBD3L4", "MBD3L5", "PRAMEF1", "PRAMEF10", "PRAMEF11", "PRAMEF12", "PRAMEF13", "PRAMEF14", "PRAMEF15", "PRAMEF17", "PRAMEF18", "PRAMEF19", "PRAMEF2", "PRAMEF3", "PRAMEF4", "PRAMEF5", "PRAMEF6", "PRAMEF7", "PRAMEF8", "PRAMEF9", "RFPL2", "RFPL4B", "RP11-432M8.11", "RP11-432M8.17", "RP11-432M8.9", "RP11-554D14.4", "RP13-221M14.1", "RP13-221M14.3", "RP13-221M14.5", "SLC34A2", "TPRX1", "TRIM43", "TRIM43B", "TRIM43CP", "TRIM49","TRIM49B", "TRIM49C", "TRIM49DP", "TRIM49L1", "TRIM51", "TRIM51BP", "TRIM51CP", "TRIM51EP", "TRIM53AP", "TRIM53BP", "TRIM53CP", "WI2-2994D6.1", "WI2-2994D6.2", "WI2-3308P17.2", "XX-FW84067D5.1", "ZNF280A", "ZNF705G", "ZSCAN4")

#vDUX4 dataset
dataset = "psmdata.geneid.static"
i <- get(dataset)
yao.biomarkers <- i[which(i$geneid %in% yao.genes), ]

yao.biomarkers <- yao.biomarkers %>% filter(rep !=3, rep !=4)
g <- ggplot(aes(y = norm.hl.ratio, x = geneid, fill = ""), data = yao.biomarkers)
g + xlab("") + ylab("log2 (DUX4/Ctrl)") +
geom_abline(intercept = 0, slope = 0, linetype = "dashed") +
geom_boxplot(width= 0.7, outlier.shape=NA) +
ylim(c(-10, 15)) +
#coord_flip() +
scale_fill_manual(name = "", values = c("#0298B820")) +
geom_jitter(height = 0, color = ifelse(yao.biomarkers$rep == 1, "#0298B8", "#810f7c"), size = 1) +
theme(panel.background = element_blank(),
panel.border = element_rect(fill=NA),
panel.grid.major.y = element_blank(),
panel.grid.minor.y = element_blank(),
text = element_text(size=10))

#iDUX4 dataset
dataset = "psmdata.geneid.static"
i <- get(dataset)
yao.biomarkers <- i[which(i$geneid %in% yao.genes), ]

yao.biomarkers <- yao.biomarkers %>% filter(rep !=1, rep !=2)
g <- ggplot(aes(y = norm.hl.ratio, x = geneid, fill = ""), data = yao.biomarkers)
g + xlab("") + ylab("log2 (DUX4/Ctrl)") +
geom_abline(intercept = 0, slope = 0, linetype = "dashed") +
geom_boxplot(width= 0.7, outlier.shape=NA) +
ylim(c(-10, 15)) +
#coord_flip() +
scale_fill_manual(name = "", values = c("#90000050")) +
geom_jitter(height = 0, color = ifelse(yao.biomarkers$rep == 3, "red", "#5e3c99"), size = 1) +
theme(panel.background = element_blank(),
panel.border = element_rect(fill=NA),
panel.grid.major.y = element_blank(),
panel.grid.minor.y = element_blank(),
text = element_text(size=10))
```
Figure 2
```{r}
#scatter plot
data <- data.rnaseq.silac.static %>% filter(rep !=3, rep != 4)

plot(data$iDUX4_logFC_UF, data$hl.ratio, pch = 21, col = "#10101040", 
     bg = "#10101020",
     xlim = c(-10, 10), ylim = c(-10, 10))
abline(h = 0, v = 0, col = "grey60", lty = 2)
```

Figure 3
```{r}
#scatter plot
data <- data.rnaseq.silac.static %>% filter(rep !=3, rep != 4)

plot(data$iDUX4_logFC_UF, data$hl.ratio, pch = 21, col = "#10101040", 
     bg = "#10101020",
     xlim = c(-3, 3), ylim = c(-3, 3))
abline(h = 0, v = 0, col = "grey60", lty = 2)

#Fig 3B panels
x <- data.rnaseq.silac.static %>% 
        select(geneid, iDUX4_logFC, vDUX4_logFC, 
              hl.ratio.rep1, hl.ratio.rep2, hl.ratio)

names(x) <- c("geneid", "5-iDUX4_mRNA_14h", "3-vDUX4_mRNA_36h", 
              "2-vDUX4_protein_24h", "1-vDUX4_protein_36h", 
              "4-iDUX4_protein_14h_median")

x <- x %>% filter(geneid == "HSPA5") %>% gather(, value, 2:6)
       
# draw the histogram with the specified number of bins
ggplot(x, aes(x = key, y = as.numeric(value))) + 
      mytheme +
      ylab("DUX4/Control ratio (log2)") +
      xlab("") +
      scale_y_continuous(limits=c(-2, 2)) +
      ggtitle("DUX4") +
      geom_bar(stat = "identity") +
      coord_flip() 
```

Figure 4
```{r}
## Post-transcriptional gene regulation
data <- data.rnaseq.silac.static %>% filter(rep !=3, rep != 4)

plot(data$iDUX4_logFC_UF, data$hl.ratio, pch = 21, col = "#10101040", 
     bg = ifelse((data$iDUX4_logFC_UF < 0.5 & data$hl.ratio > 1), "gold", 
                 ifelse((data$iDUX4_logFC_UF > -0.5 & data$hl.ratio < -1), "gold", "#10101040")),
     xlim = c(-5, 10), ylim = c(-10, 10))
abline(h = 0, v = 0, col = "grey60", lty = 2)

#NMD factor plot
load("data/rdata/uniprot.ids.sequences.Rdata")
gene.annotation <- uniprot.ids.sequences[, c(2, 3)]
nmd.genes <- c("UPF1", "UPF2", "UPF3B", "SMG1", "SMG5", "SMG6", "SMG7", "DCP1A", "DCP2", "XRN1", "ETF1", "GSPT1", "GSPT2", "RBM8A", "MAGOH", "CASC3", "EIF4A3")
nmd.genes <- data.frame(unique(nmd.genes))
names(nmd.genes) <- c("geneid")

tmp <- inner_join(nmd.genes, data.rnaseq.silac.static, by = "geneid")
tmp <- tmp %>% filter(!is.na(iDUX4_logFC_UF), !is.na(hl.ratio))
tmp <- tmp[, c(1, 2, 38)]

plot(tmp$iDUX4_logFC_UF, tmp$hl.ratio, 
     pch = 21, col = "black", 
     bg = "red", 
     xlim = c(-5, 10), 
     ylim = c(-5, 10))
abline(h = 0, v = 0, col = "grey60", lty = 2)

#ubiquitin ligase plot
library(readr)
genes <- read_delim("analysis/geneset.txt", 
    "\t", escape_double = FALSE, col_names = FALSE, 
    trim_ws = TRUE)
names(genes) <- c("geneid")

load("data/rdata/uniprot.ids.sequences.Rdata")
gene.annotation <- uniprot.ids.sequences[, c(2, 3)]
ubiquitin.genes1 <- gene.annotation %>% filter(grepl('ubiquitin', gene))
ubiquitin.genes2 <- gene.annotation %>% filter(grepl('proteasome', gene))
ubiquitin.genes3 <- gene.annotation %>% filter(grepl('TRIM', geneid))
ubiquitin.genes4 <- gene.annotation %>% filter(grepl('RFPL', geneid))

ubiquitin.genes <- rbind(ubiquitin.genes1, ubiquitin.genes2, ubiquitin.genes3, ubiquitin.genes4)
ubiquitin.genes <- ubiquitin.genes[, c(1)]
ubiquitin.genes <- data.frame(unique(ubiquitin.genes))
names(ubiquitin.genes) <- c("geneid")

ubiquitin.genes.master <- rbind(ubiquitin.genes, genes)
ubiquitin.genes.master <- unique(ubiquitin.genes.master)
tmp <- inner_join(ubiquitin.genes.master, data.rnaseq.silac.static, by = "geneid")
tmp <- tmp %>% filter(!is.na(iDUX4_logFC_UF), !is.na(hl.ratio))

tmp <- tmp %>% filter(rep !=3, rep != 4)
plot(tmp$iDUX4_logFC_UF, tmp$hl.ratio, 
     pch = 21, col = "black", 
     bg = "red", 
     xlim = c(-5, 10), 
     ylim = c(-5, 10))
abline(h = 0, v = 0, col = "grey60", lty = 2)

identify(tmp$iDUX4_logFC_UF, tmp$hl.ratio, labels=tmp$geneid, cex = 1)
```
````
